# Unraveling the molecular mechanism underlying the anticancer activity of CISD2/NAF-1^44-67^

**DOI:** 10.1101/2025.01.07.630758

**Authors:** Linda Rowland, Itai Alfoni, Ehud Neumann, Ola Karmi, Rachel Nechushtai, Ron Mittler

## Abstract

We recently reported on the development of a unique cancer-targeting peptide called NAF-1^44-67^ (derived from CISD2/NAF-1). NAF-1^44-67^ selectively permeates the plasma membrane (PM) of cancer cells, but not healthy cells, causing the activation of apoptotic and ferroptotic cell death pathways specifically in cancer cells. NAF-1^44-67^ also targets and shrinks human breast and ovarian cancer tumors in a xenograft mice model system without any apparent side effects. Although the specific permeation of NAF-1^44-67^ through cancer cell PMs was studied, and its cancer killing effects validated *in vitro* and *in vivo*, little is known about how NAF-1^44-67^ exerts its biological activity once it enters cancer cells. Here, we report that NAF-1^44-67^ targets the CISD2/NAF-1 protein of cancer cells and disrupts its homodimeric structure. We further reveal that a peptide derived from the same domain of the human CISD1 (mitoNEET; mNT^19-42^) protein, a close family member to CISD2, has no killing activity towards cancer cells, and that dimers of NAF-1^44-67^ (at two different orientations) have higher anticancer activity compared to monomeric NAF-1^44-67^. Our findings shed new light on the biological activity of NAF-1^44-67^ and bring it closer to becoming a potential new anticancer drug.

## Introduction

The CDGSH iron sulfur domain 2 (CISD2) protein plays a key role in regulating cancer cell proliferation, metastasis, and survival, and is used as a prognostic marker for several different cancers (Table S1). CISD2 is a mitochondria and endoplasmic reticulum (ER) membrane-anchored NEET protein (also called Nutrient Autophagy Factor 1; NAF-1) that interacts with several different proteins involved in calcium signaling and cell death regulation [1,2]. A prominent role of CISD2/NAF-1 in cancer cells is to prevent the activation of ferroptosis and apoptosis by keeping the levels of free iron and iron-sulfur (Fe-S) clusters under control, preventing the overaccumulation of reactive oxygen species (ROS) [2-5].

We recently reported on the discovery of a unique cancer-targeting peptide (CTP) termed NAF-1^44-67^ [6-8]. This peptide is derived from part of the transmembrane domain of CISD2/NAF-1, as well as the unstructured domain that is immediately adjacent to it (Fig. 1A). NAF-1^44-67^ was found to selectively kill several different epithelial breast cancer cell lines, without affecting control epithelial cells [6,7]. It selectively permeates the plasma membrane (PM) of human epithelial breast cancer cells (MDA-MB-231), but not control epithelial cells (MCF-10A), causing the activation of apoptotic and ferroptotic cell death pathways, specifically in cancer cells [6,7]. Recently, we reported that NAF-1^44-67^ can also specifically target human ovarian cancer cells, as well as multiple other human cancer cell lines [8]. In addition, we showed that NAF-1^44-67^ targets and shrinks human breast and ovarian cancer tumors in a xenograft mice model system without any apparent side effects [6,8].

**Fig. 1.**
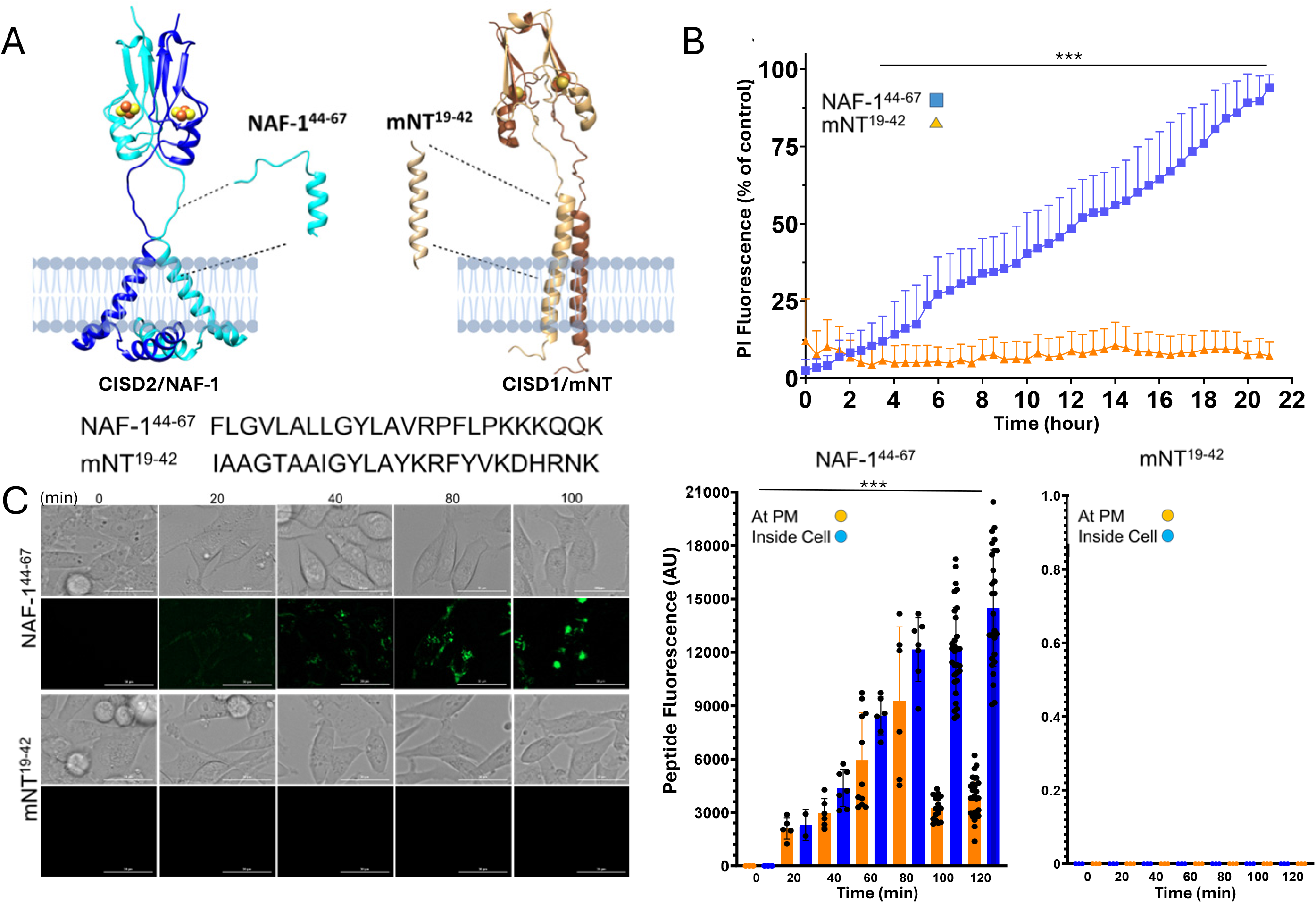
Unlike NAF-1^44-67^, derived from CISD2, mNT^19-42^, derived from CISD1, does not have anticancer killing activity. **(A)** A model showing the origins, sequence, and predicted structures of NAF-1^44-67^ and mNT^19-42^. **(B)** Cell viability measurements of MDA-MB-231 human breast cancer cells treated with 15μM NAF-1^44-67^ or mNT^19-42^. **(C)** Representative images (left; bar is 30 μ m) and bar graphs quantifying peptide fluorescence at the plasma membrane or inside cells (right) of MDA-MB-231 cells treated with 15μM of fluorescently [5(6)-carboxyfluorescein]-labelled NAF-1^44-67^ or mNT^19-42^. All experiments were repeated with 3 biological repeats, each with at least 3 technical repeats. Statistical significance was determined using two-tailed student’s t-test or one-way ANOVA (*** *p* < 0.01). Abbreviations: AU, arbitrary units; mNT, mitoNEET; NAF-1, nutrient autophagy factor 1; PI, propidium iodide.

Although the specific permeation of NAF-1^44-67^ through cancer cell PMs was studied, and its cancer killing effects validated *in vitro* and *in vivo* [6-9], little is known about how NAF-1^44-67^ exerts its biological activity once it enters cancer cells. In addition, it is unknown whether other peptides, derived from other CISD/NEET proteins will have similar activity to that of NAF-1^44-67^, as well as whether aggregation of NAF-1^44-67^ at the surface of the cancer PM will enhance its anticancer activity. Here we report that a peptide derived from the same domain of the human CISD1 (mitoNEET; mNT^19-42^) protein, a close family member to CISD2 [2,10], has no killing activity towards human breast cancer cells. We further reveal that unlike NAF-1^44-67^, mNT^19-42^ does not bind to and permeates human breast cancer cells. Using dimers of NAF-1^44-67^ (at two different orientations) we further show that the anticancer activity of NAF-1^44-67^ is enhanced upon its dimerization. Finally, using a split-yellow fluorescence protein (split-YFP) approach we demonstrate that NAF-1^44-67^ targets the CISD2/NAF-1 protein of cancer cells and disrupts its homodimeric structure. Our findings shed new light on the biological activity of NAF-1^44-67^ and bring it closer to becoming a potential new anticancer drug.

## Materials and Methods

Cell lines were obtained and grown as described in [7]. Peptides and other materials were obtained as described in [6-8]. Cell death and split-YFP measurements were conducted as described in [7,11]. Structures were obtained from [https://www.rcsb.org/] or generated using alphafold (https://alphafoldserver.com). Statistical analysis was conducted according to [6,7].

## Results and Discussion

### Specificity in the anticancer killing activity of NAF-1^44-67^

To determine whether the anticancer killing activity of NAF-1^44-67^ is unique to the protein it originated from (CISD2/NAF-1), we synthesized a peptide derived from the same domain of its closest mammalian homolog, CISD1/mNT (Fig. 1A). CISD1/mNT and CISD2/NAF-1 share 54% identical residues and 69% similar residues, and together participate in the same pathway that regulates the transfer of 2Fe-2S clusters from within the mitochondria to the cytosol, essential for cancer cell survival [2-5,10-12]. Compared to the killing effect of NAF-1^44-67^ towards MDA-MB-231 cells, mNT^19-42^ did not display a significant anticancer activity (Fig. 1B). This finding suggests that the anticancer killing activity of NAF-1^44-67^ is dependent on the protein it originated from. It is also in agreement with a previous study showing that suppressing CISD2/NAF-1 expression in cancer cells has a much stronger inhibitory effect on cancer cell proliferation and xenograft tumor growth, compared to suppressing CISD1/mNT expression [3].

The low anticancer activity of mNT^19-42^ could result from its inability to permeate cancer cells or from its biological activity once it entered cancer cells. We therefore compared between the cell permeating activity of the two peptides. Compared to NAF-1^44-67^ that binds to the PMs of MDA-MB-231 cells and permeates them, mNT^19-42^ did not bind to, or permeated, breast cancer cells (Fig. 1C). This finding reveals that mNT^19-42^ does not share the unique features of NAF-1^44-67^ that allows it to bind and permeate the PM of cancer cells [6-9]. As the two peptides share 33% identical residues and 46% similar residues and both have a charged C-terminal (5 charged residues in NAF-1^44-67^ and 6 in mNT^19-42^; Fig. 1A), further studies are needed to determine what distinguishes NAF-1^44-67^ from mNT^19-42^ and allow it to selectively kill cancer cells.

### Dimerization of NAF-1^44-67^ enhances its anticancer killing activity

As demonstrated in Fig. 1, as well as reported previously [6-9], the cancer PM-specific permeating activity of NAF-1^44-67^ is highly intriguing. Computational modeling of this process suggested that NAF-1^44-67^ first binds to the cancer PM via its negative charge and then permeates it due to its amphipathic properties [6,9]. However, whether the peptide aggregates on the surface of the cancer cell PM, like some anticancer peptides do [13,14], is unknown. To address this question, we synthesized two dimers of NAF-1^44- 67^ in two different orientations (Fig. 2A; bound together via their N- or C-terminals; Dimers 1 or 2, respectively). We then compared the anticancer killing activity of the two NAF-1^44-67^ dimers to that of the NAF-1^44-67^ monomer. Compared to the killing effect of the NAF-1^44-67^ monomer, both NAF-1^44-67^ dimers had a higher anticancer activity (Fig. 2A). This finding suggests that aggregation of NAF-1^44-67^ could help it permeate cancer cells. Increasing the concentration/dose of NAF-1^44-67^ was previously shown to increase its anticancer activity [7] and this could happen due to increased concentration and/or aggregation of the peptide at the cancer cell PM. Alternatively, the higher toxicity of the dimer peptides may have to do with their biological activity once they entered cancer cells. As the CISD2/NAF-1 protein is a homodimer (Fig. 1A), having a dimer of the NAF-1^44-67^ peptide could make it more efficient in targeting this protein. Further studies are needed to resolve these intriguing questions. However, as monomers of NAF-1^44-67^ are efficient in killing multiple cancer cell lines, as well as targeting tumors [6-8], synthesizing the more expensive dimers might not be needed to increase toxicity; that could be achieved by simply increasing the dose of the monomer peptide.

**Fig. 2.**
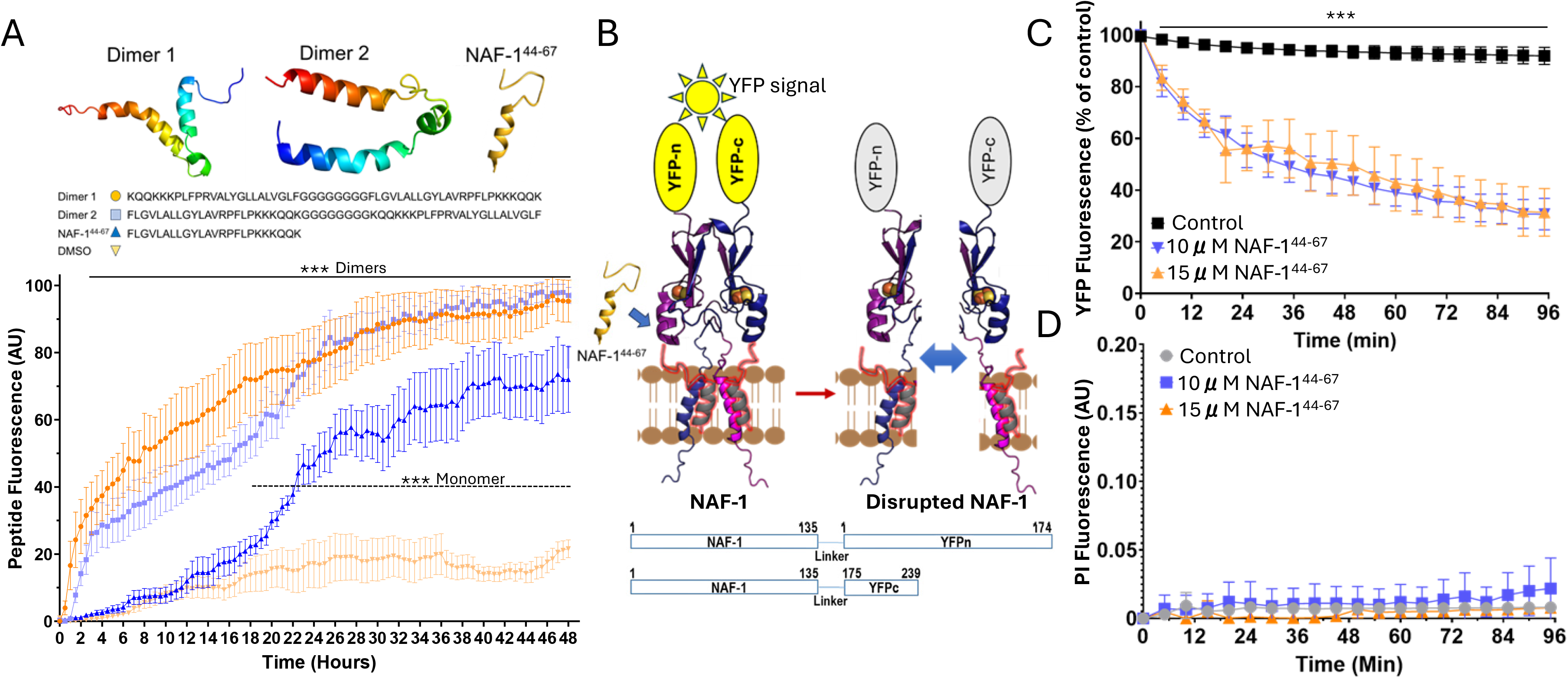
NAF-1^44-67^ targets the CISD2 protein of cancer cells, and its dimerization enhances its toxicity. **(A)** Predicted structures and sequences (top), and a graph showing cell viability measurements of MDA-MB-231 breast cancer cells treated with 15μM of NAF-1^44-67^ Dimer 1, NAF-1^44-67^ Dimer 2, or the NAF-1^44-67^ monomer (bottom). **(B)** A model showing the split-YFP strategy used to determine whether CISD2 is a target of NAF-1^44-67^ in cancer cells. **(C)** YFP fluorescence of MDA-MB-231 cells expressing *NAF-1*^*1-135*^*-YFPc* and *NAF-1*^*1-135*^*-YFPn* in the presence or absence of 10 or 15 μM NAF-1^44-67^. (D) Cell viability measurements of MDA-MB-231 cells expressing *NAF-1*^*1-135*^*-YFPc* and *NAF-1*^*1-135*^*-YFPn* in the presence or absence of 10 or 15 μM NAF-1^44-67^. All experiments were repeated with 3 biological repeats, each with at least 3 technical repeats. Statistical significance was determined using two-tailed student’s t-test or one-way ANOVA (*** *p* < 0.01). Abbreviations: AU, arbitrary units; NAF-1, nutrient autophagy factor 1; PI, propidium iodide; YFP, yellow fluorescence protein.

### NAF-1^44-67^ targets the CISD2/NAF-1 protein of cancer cells

NAF-1^44-67^ is derived from the CISD2/NAF-1 homodimeric protein and shows specificity to this protein (Fig. 1). In addition, dimers of NAF-1^44-67^, containing two copies of its partial membrane spanning domain, are more toxic compared to monomeric NAF-1^44-67^ (Fig. 2A). We therefore hypothesized that once NAF-1^44-67^ enters cancer cells it could target their CISD2/NAF-1 protein and disrupt its homodimeric structure (Fig. 2B). To test this possibility, we used MDA-MB-231 cells that express subunits of the homodimeric CISD2/NAF-1 protein with two different halves of the YFP protein attached in frame to each of their C-terminals (*NAF-1*^*1-135*^*-YFPc* and *NAF-1*^*1-135*^*-YFPn*; Fig. 2B). These cells were developed as controls for a study that focused on the protein-protein interaction between CISD2/NAF-1 and CISD1/mNT and display a strong YFP signal in cells grown in culture [11]. Treatment of MDA-MB-231 cells that display the CISD2/NAF-1 homodimer-generated YFP signal with NAF-1^44-67^ caused the signal to quench within minutes (Fig. 2C). To ascertain that within this time frame (0-96 min) no cell death (that could also cause the YFP signal to quench) occurs, we also measured cell viability of the *NAF-1*^*1-135*^*-YFPc* and *NAF-1*^*1-135*^*-YFPn* MDA-MB-231 cells treated with NAF-1^44-67^. In agreement with the results shown in Figs. 1B, and 2A, no cell death was detected within this time frame in MDA-MB-231 cells treated NAF-1^44-67^ (Fig. 4D). As previously reported, the integrity of mitochondria and ER of MDA-MB-231 cells, to which CISD2/NAF-1 is localized [2,11], is also not affected within this time frame by NAF-1^44-67^ [6]. Taken together, our results suggest that the CISD2/NAF-1 protein of cancer cells is one of the targets of NAF-1^44-67^ and that the peptide disrupts its homodimeric structure.

### Summary

CISD2/NAF-1 plays a key role in maintaining the levels of ROS, iron, and Fe-S clusters under control in cancer cells [2-5,11,12,15]. Any disruption in its activity or a decrease in its protein level were shown to cause the activation of ferroptosis and apoptosis in cancer cells (Table S1). We previously demonstrated that expressing a mutated subunit of CISD2/NAF-1 (H114C; unable to donate this protein’s 2Fe-2S clusters to acceptor proteins), in cancer cells causes a dominant negative effect on CISD2/NAF-1 function triggering cancer cell death [4,15]. These findings support a model in which CISD2/NAF-1 needs to maintain its homodimeric structure to function in cancer cells. NAF-1^44-67^ that specifically permeates cancer cells and disrupts the homodimeric structure of their CISD2/NAF-1 protein (Figs. 1, 2) could therefore trigger ferroptotic and apoptotic cells death in these cells. Proteomics of MDA-MB-231 cells treated with NAF-1^44-67^ has indeed revealed that both cell death pathways are activated in cancer cells following treatment with the peptide [6]. Taken together, our work suggests that NAF-1^44-67^ could be used as a cancer-specific drug that targets the dependency of cancer cells on high iron and ROS levels, required for proliferation and metastasis, via targeting of the CISD2/NAF-1 protein.

## Authorship contribution statement

Linda Rowland: Methodology, Formal analysis, Data curation; Itai Alfoni, Ehud Neumann, and Ola Karmi: Formal analysis, Data curation; Rachel Nechushtai and Ron Mittler: Writing, review and editing, Supervision, Project administration, Funding acquisition, Conceptualization.

## Consent for publication

Not applicable.

## Availability of data and material

This published article and its supplementary information files include all data generated or analyzed during this study.

## Funding

This work was supported by the National Institute of Health grant GM111364 (to R.M.), the National Science Foundation (NSF)-Binational Science Foundation (BSF) Grant NSF-MCB 1613462 (to R.M.) and BSF Grant 2015831 (to R.N.).

## Declaration of competing interest

The authors declare no conflict of interest.

## Institutional review or ethical board

Not applicable.

---

**Table S1.**
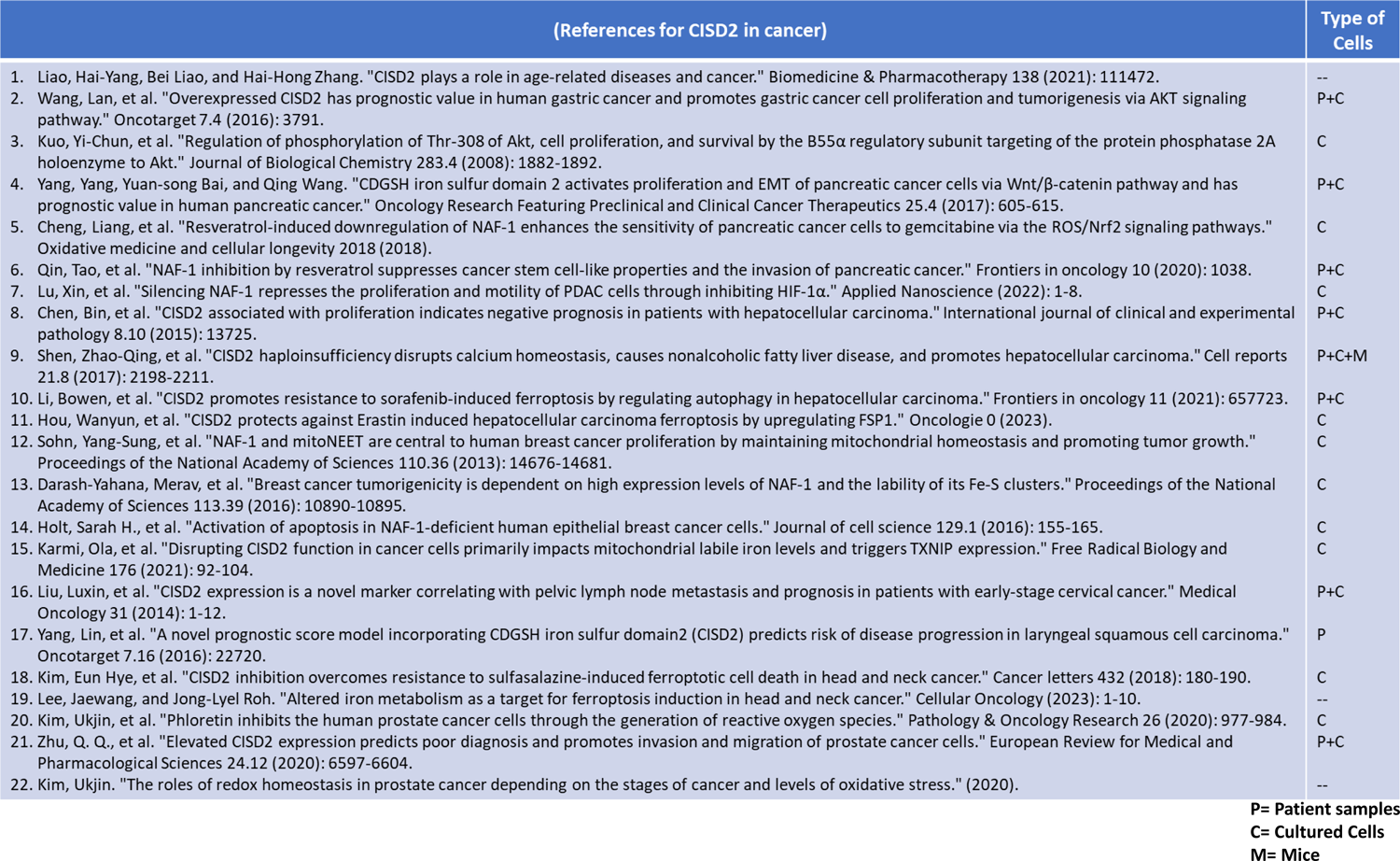
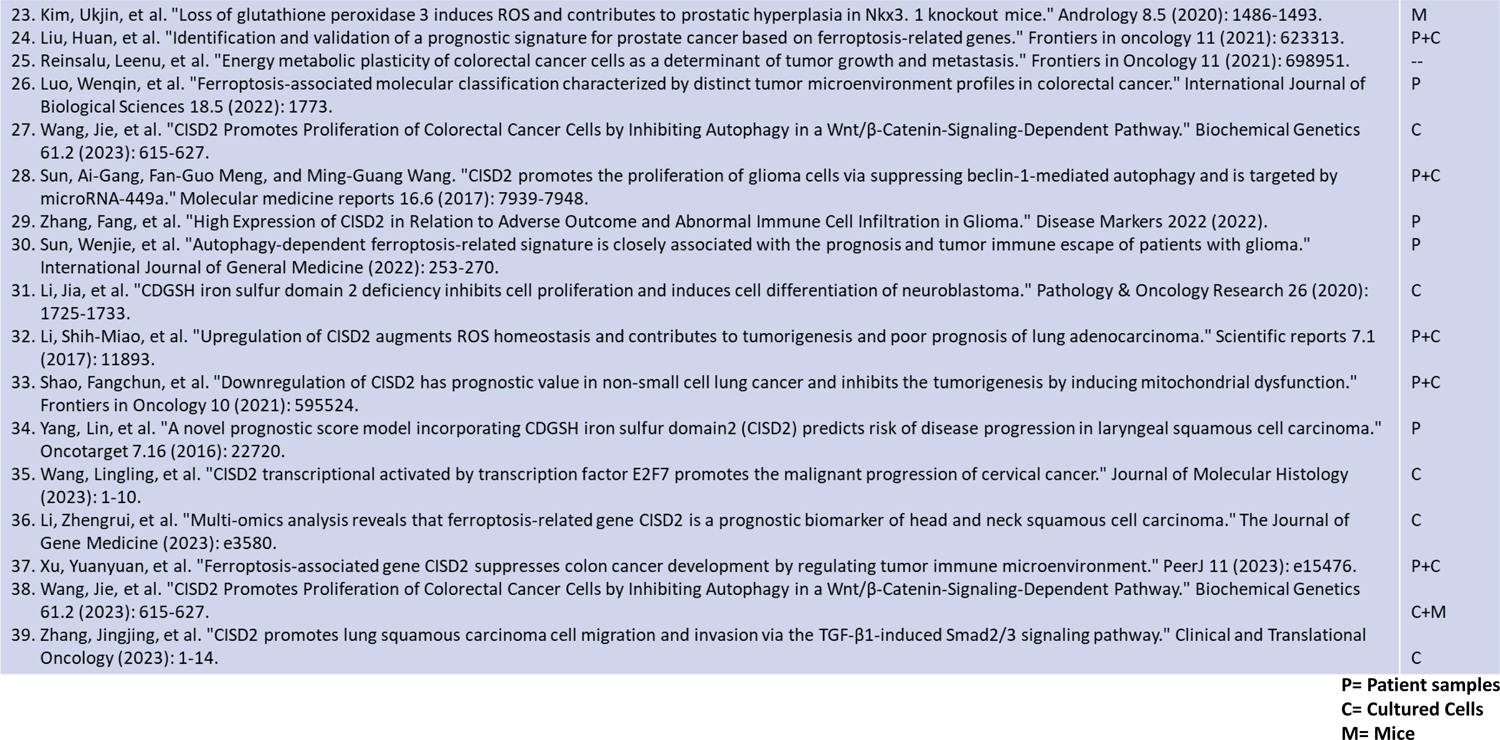
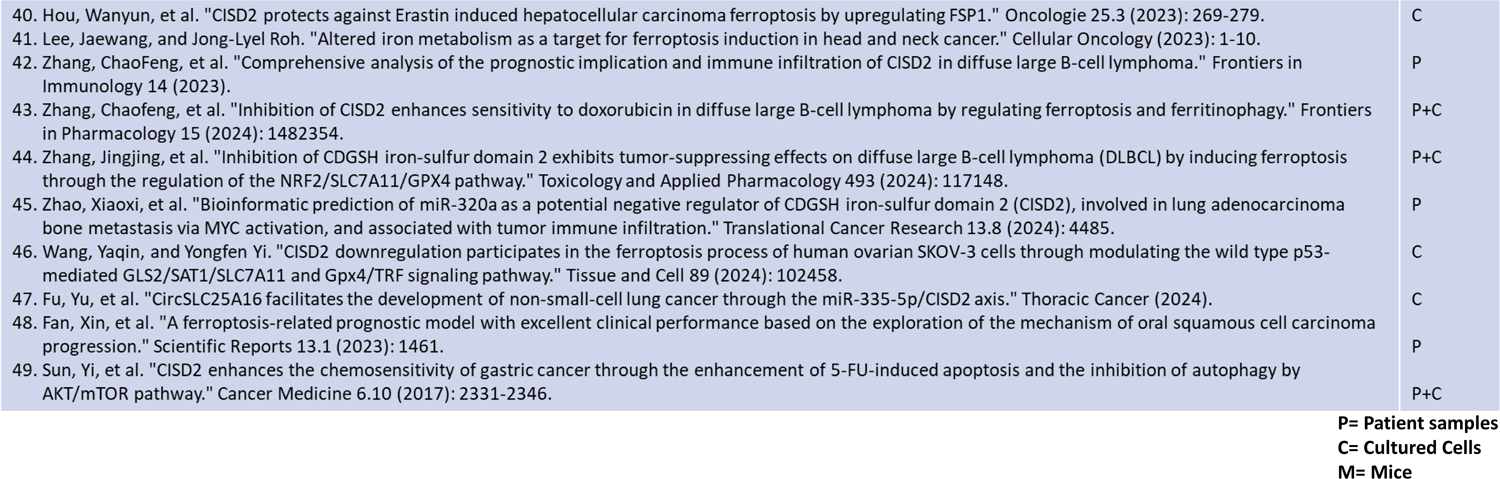
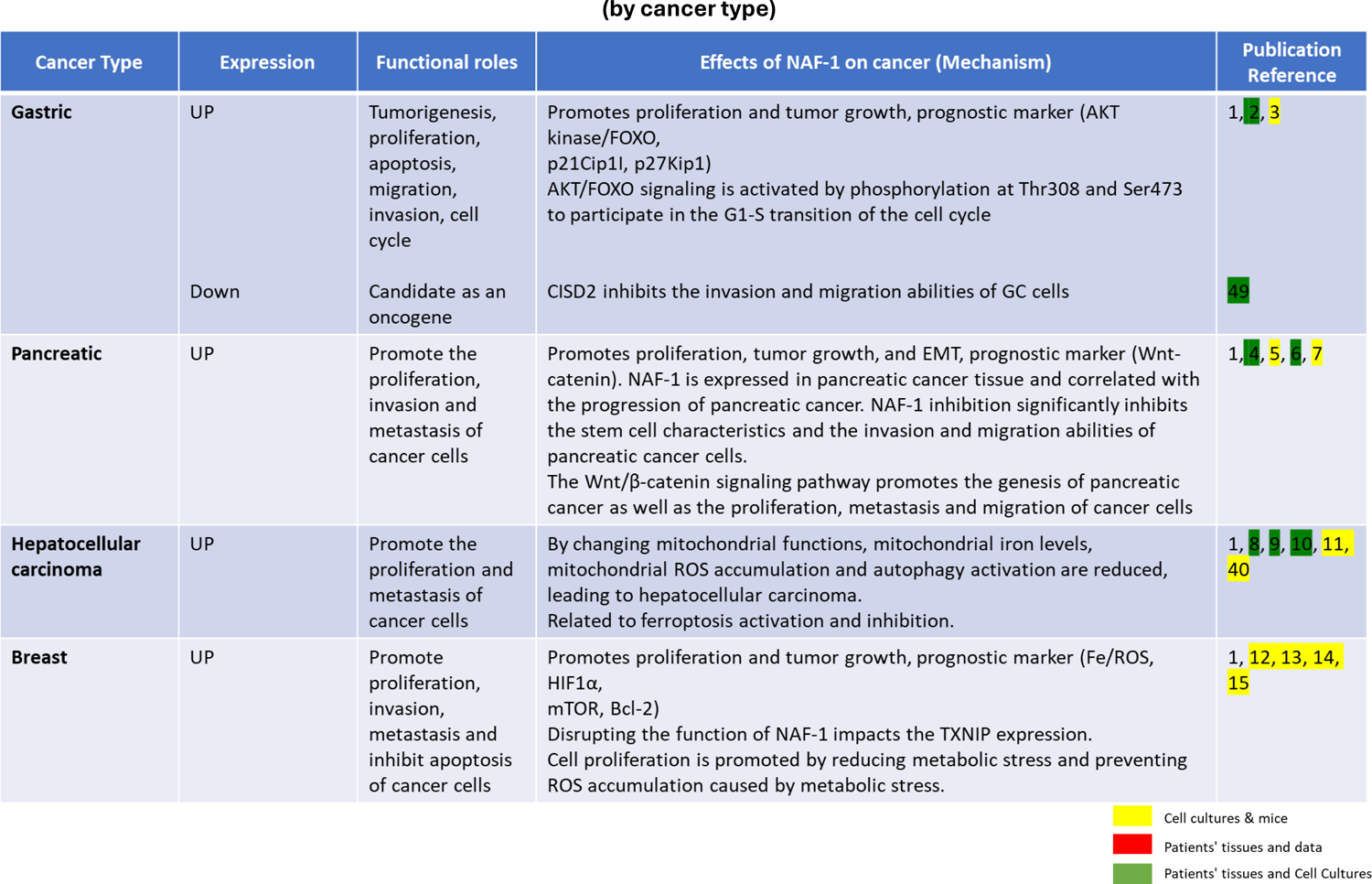
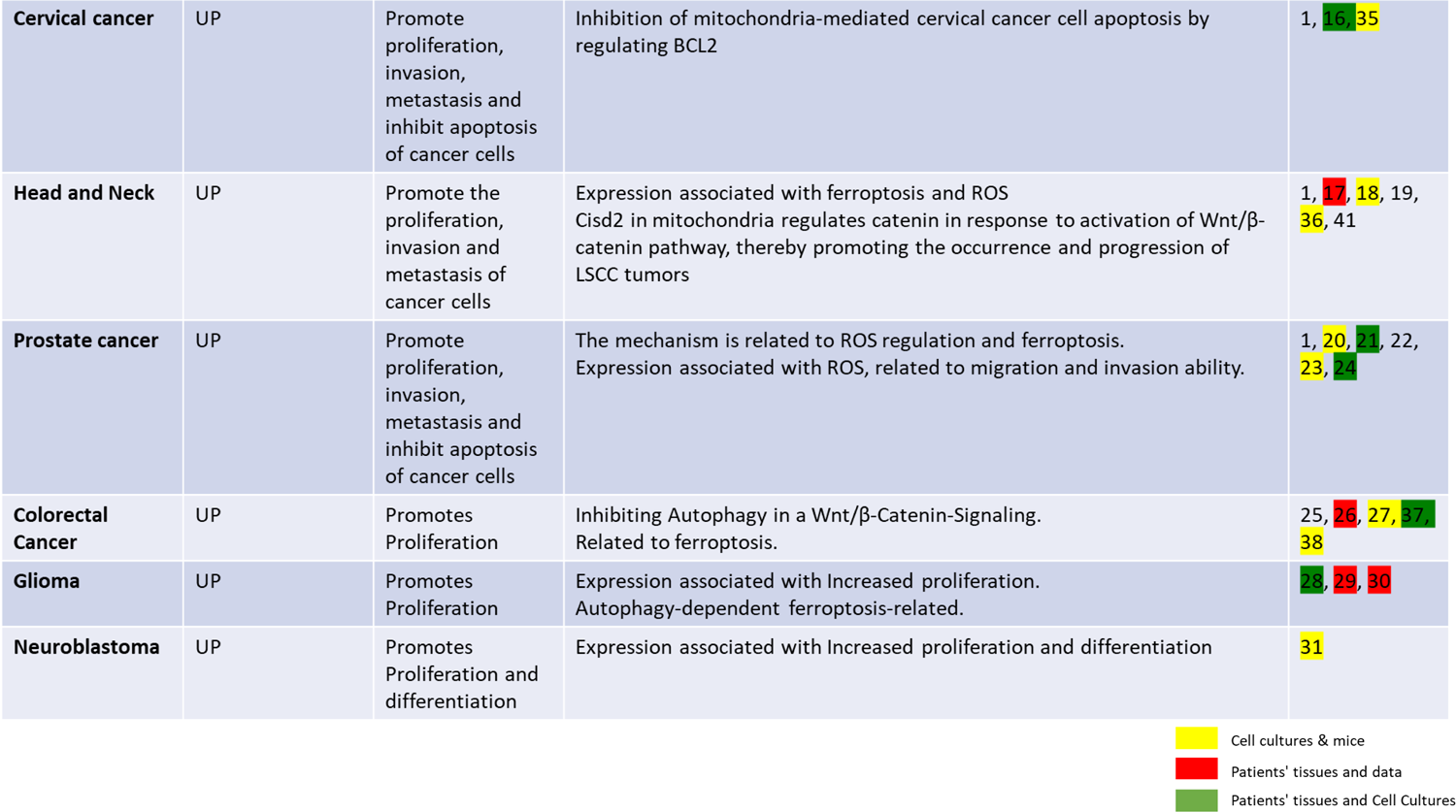
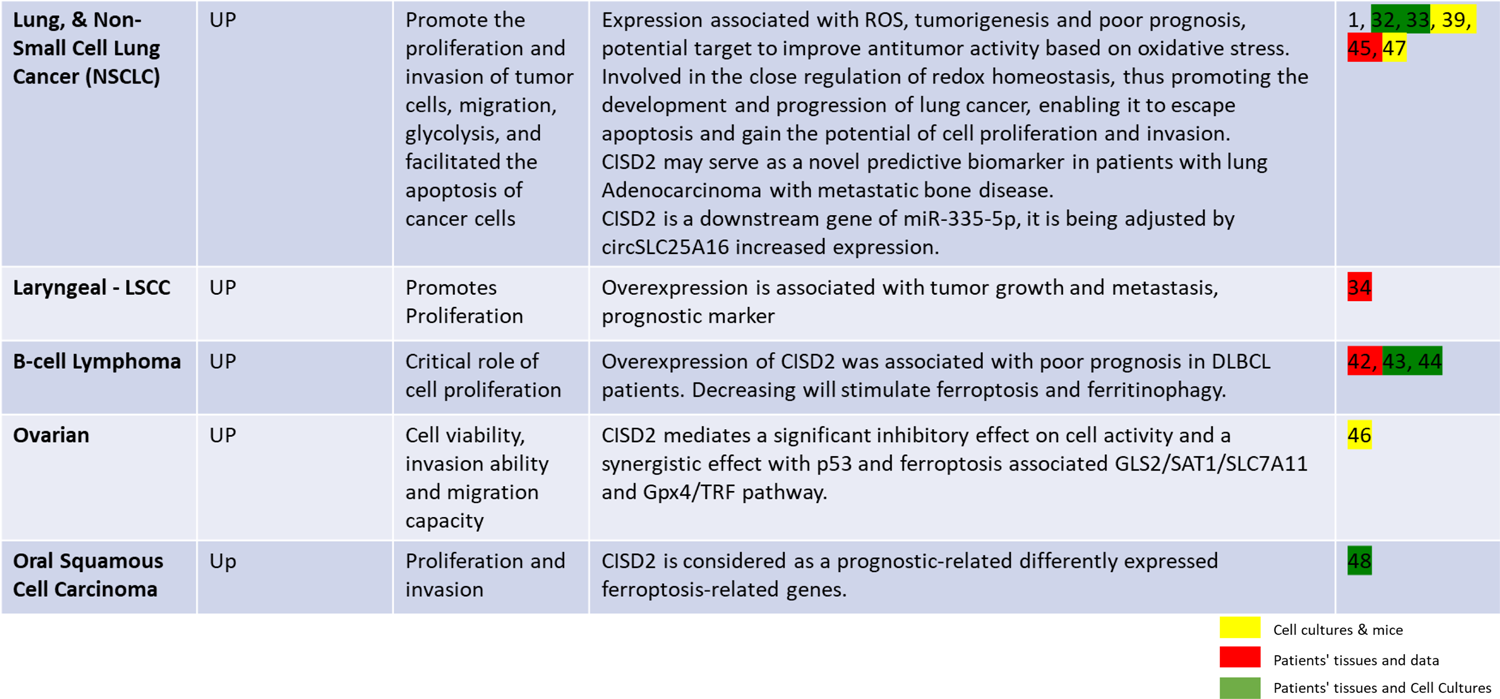

## Notes

### Competing Interest Statement

The authors have declared no competing interest.

